# A universal resazurin-based viability assay for prokaryotic and eukaryotic cells in 2D and 3D cultures

**DOI:** 10.64898/2026.04.13.718248

**Authors:** Ramón Cervantes-Rivera, Atalia Ziret Romero Rosas, Sandra Jetsamari Figueroa Ortíz, Luisa Nirvana González-Fernández, Alejandra Ochoa-Zarzosa, Joel E. López-Meza

## Abstract

*In vitro* cytotoxicity assessments frequently rely on staining-based methods that indirectly estimate viable cell numbers indirectly. A major limitation of many such techniques is their endpoint nature, requiring cell lysis or irreversible processing that precludes longitudinal monitoring of cellular responses following treatment. An ideal assay for evaluating cell viability and proliferation should be simple, rapid, cost-effective, reproducible, and highly sensitive, while also enabling accurate quantification with minimal interference from test compounds.

The resazurin reduction assay satisfies these criteria, offering a sensitive and economical alternative to conventional tetrazolium-based methods. Although both assay types depend on the metabolic reduction of a dye by viable cells, they differ mechanistically. Tetrazolium salts (e.g., MTT) are reduced by cellular dehydrogenases to insoluble formazan crystals that require solubilization before to detection. In contrast, resazurin—a cell-permeable, non-fluorescent blue dye—is reduced to resorufin, a highly fluorescent compound detectable without additional processing steps. This property renders the resazurin assay broadly applicable to viability testing in eukaryotic cells cultured in both 2D and 3D formats, as well as in bacterial systems.

Here, we present a streamlined, universal protocol for implementing the resazurin reduction assay across diverse experimental models, emphasizing its practicality, reproducibility, and adaptability for real-time viability monitoring.

**Key features:**

- Real-time, non-destructive monitoring: Enables longitudinal studies by allowing repeated measurements of the same samples over hours without toxicity or disruption.
- Streamlined workflow: A simple “add-incubate-read” protocol eliminates the need for cell lysis, washing, or extraction, saving time and reducing variability.
- Broad sample compatibility: Versatile and reliable for use with 2D monolayers, 3D spheroids, organoids, and bacterial cultures.
- High sensitivity: Fluorescent detection of resorufin provides exceptional sensitivity, enabling accurate quantification of even small viable cell populations.
- Low background and minimal interference: A clean fluorescent readout reduces the risk of signal artifacts, offering a more reliable alternative to traditional colorimetric assays.
- Cost-effective and accessible: Utilizes standard laboratory plate readers and commercially available reagents, making it an economical choice for any lab.
- Scalable for high-throughput screening: Easily adaptable to various plate formats, supporting both small-scale experiments and large-scale automated screening applications.

**Graphical overview:** 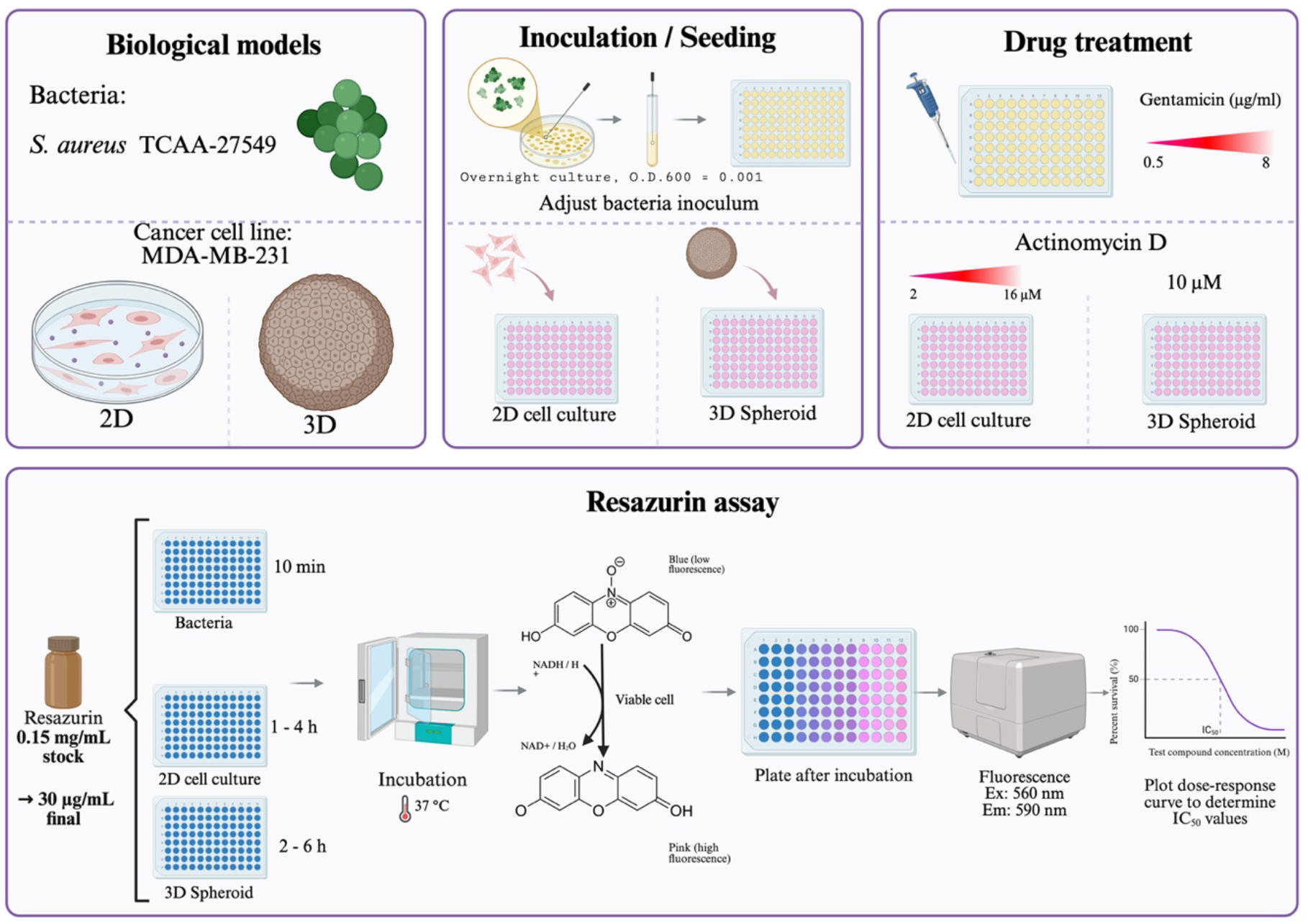

## Background

The assessment of cell viability and metabolic activity constitutes a fundamental pillar of biomedical research, underpinning applications ranging from basic toxicology and pharmacology to high-throughput drug screening, tissue engineering, and microbiological quality control(Njoku et al., 2023; Vieira-da-Silva & Castanho, 2023). As the complexity of experimental models has evolved from traditional two-dimensional (2D) monolayers to three-dimensional (3D) spheroids, organoids, and co-culture systems, so too has the demand for assay methods that are not only accurate and reproducible but also simple, cost-effective, non-toxic, and amenable to longitudinal study designs(Lavogina et al., 2022; Njoku et al., 2023).

Resazurin is a cell-permeable, low-toxicity dye that exhibits minimal fluorescence in its oxidized state(Njoku et al., 2023). Upon entering viable cells, it is reduced by metabolic enzymes, utilizing electron donors such as NADPH, FADH, and FMNH. The product, resorufin, is released into the culture medium and can be detected quantitatively using either fluorometric or colorimetric methods, although fluorescence detection offers superior sensitivity. The quantity of resorufin generated is directly proportional to the number of metabolically active cells under optimized conditions, enabling robust estimation of viable cell populations(Vieira-da-Silva & Castanho, 2023).

Before the widespread adoption of resazurin, tetrazolium salts—most notably MTT (3-(4,5-dimethylthiazol-2-yl)-2,5-diphenyltetrazolium bromide)—represented the dominant methodology for viability assessment. Both assay classes rely on metabolic reduction by viable cells, yet they differ in several practically significant respects. Tetrazolium compounds are reduced to insoluble formazan crystals that require solubilization in organic solvents before spectrophotometric quantification—an additional processing step that increases assay time, introduces variability, and precludes the return of live cells to culture. Resazurin, by contrast, yields a soluble, intrinsically fluorescent product that is directly detectable in the culture medium without cell lysis, washing, or extraction steps(Emter & Natsch, 2015). This non-destructive character represents the assay’s cardinal advantage: the same cell population can be monitored repeatedly over hours, enabling genuine longitudinal studies of proliferation, cytotoxicity, and recovery from treatment(Vieira-da-Silva & Castanho, 2023) . Furthermore, resazurin assays demonstrate comparable or superior sensitivity to MTT, with lower inter-assay variability and enhanced compatibility with high-throughput screening formats(Emter & Natsch, 2015).

The versatility of the resazurin reduction assay is reflected in the extraordinary range of biological systems to which it has been successfully applied. In eukaryotic research, it supports viability assessment in immortalized cell lines, primary cells, and stem cells cultured in both conventional 2D monolayers and complex 3D architectures such as spheroids and scaffold-based constructs. In microbiology, it enables rapid antimicrobial susceptibility testing and quantification of bacterial and fungal viability. This cross-kingdom applicability—uncommon among viability assays—derives from the fundamental conservation of cellular redox metabolism(Njoku et al., 2023).

Despite its widespread use, the resazurin assay is not without limitations that warrant careful consideration. First, it is essential to recognize that resazurin reduction reports metabolic activity rather than cell viability per se; cells with severely depressed metabolism may be erroneously classified as non-viable, while non-cellular reducing agents can generate false-positive signals(Vieira-da-Silva & Castanho, 2023). Second, the assay is subject to kinetic complexities: resorufin itself is a substrate for further reduction to the non-fluorescent dihydroresorufin, which can compromise linearity and lead to underestimation of viable cell numbers at extended incubation times(Vieira-da-Silva & Castanho, 2023). Third, resazurin exhibits concentration-dependent cytotoxicity in certain cell types, necessitating careful optimization of working concentrations and exposure durations(Vieira-da-Silva & Castanho, 2023). Fourth, fluorescence measurements are instrument-dependent and susceptible to inner filter effects, requiring appropriate calibration and validation. Finally, recent reviews have highlighted concerning inconsistencies in published resazurin-based studies, often attributable to poorly optimized or insufficiently standardized protocols.

The convergence of several factors—the pressing need for non-destructive, longitudinal viability assays; the demonstrated utility of resazurin across prokaryotic and eukaryotic systems in both 2D and 3D formats; and the growing recognition of reproducibility challenges arising from methodological heterogeneity—underscores the value of a standardized, universally applicable protocol. While excellent guidelines exist for specific applications or cell types, a consolidated procedure that addresses common optimization parameters (dye concentration, incubation time, detection settings) and provides practical solutions to recurrent pitfalls (kinetic nonlinearity, background interference, cytotoxicity concerns) remains conspicuously absent from the literature.

Here, we present a comprehensive, universal resazurin-based viability assay protocol optimized for use across prokaryotic and eukaryotic cells cultured in both 2D and 3D configurations. The protocol emphasizes critical control points, provides decision frameworks for assay design, and incorporates recently recommended practices for kinetic fluorescence monitoring and ratio-based calculations to ensure accurate, reproducible quantification of cellular metabolic activity. By unifying best practices from disparate application domains, this protocol serves as a practical reference for investigators seeking a reliable, adaptable, and methodologically sound approach to resazurin-based viability assessment.

## Materials and reagents

### Biological materials

1. MDA-MB-231 cell line (ATCC, catalog number: CRM-HTB-26)
2. Staphylococcus aureus (ATCC, catalog number: 27543)

### Reagents

1. Dulbecco’s modified Eagle medium (DMEM) (Sigma Merck, catalog number: D5648) or RMPI media.
2. Sodium bicarbonate (NaHCO_3_) (J.T Baker, catalog number: 3506-01)
3. L-glutamine (Gibco, catalog number: 20530-081)
4. Amphotericin B (Sigma-Aldrich, catalog number: A2942)
5. Fetal bovine serum (FBS, BioWest, catalog number: BIO-S1400)
6. Bovine calf serum (BCS, BioWest, catalog number: S0400-500)
7. Penicillin–streptomycin (Gibco, catalog number: 15140-122)
8. Trypsin (Sigma-Aldrich, catalog number: T4799-5G) (see Recipes)
9. Ethylenediaminetetraacetic acid (EDTA) (J.T Baker, catalog number: 8993-01)
10. Resazurin sodium salt (Sigma-Aldrich, catalog number: R7017)
11. Dimethyl sulfoxide (DMSO) (Sigma-Aldrich, catalog number: D2650)
12. Actinomycin D (Sigma-Aldrich, catalog number: A9415)
13. Gentamicin sulfate (SON’S, catalog number: 87882 SSA IV)
14. Trypticase soy broth (TSB) (BD Bioxon, catalog number: 211825)
15. Sodium chloride (J.T. Baker, catalog number: 3624-01)
16. Sodium phosphate, 7-hydrate, crystal (J.T. Baker, catalog number: 3824-01)
17. Monobasic potassium phosphate crystal (J.T. Baker, catalog number: 3824-01)
18. Potassium chloride (J.T. Baker, catalog number: 3040-01)
19. Sodium bicarbonate (NaHCO_3_) (J.T Baker, catalog number: 3506-01)
20. Sodium phosphate (NaH_2_PO_4_) (J.T Baker, catalog number: 3818-01)
21. Monopotassium phosphate (KH_2_PO_4_) (J.T Baker, catalog number: 3246-01)
22. Agarose (Invitrogen, catalog number: 16500-100)
23. MilliQ water sterile water (J.T. Baker, catalog number: 4220-20)

### Laboratory supplies

1. 96-well round-bottom assay plate (Corning, catalog number: 3797)
2. Non-adherent 96-well plates
3. Cell culture plate, 96 wells, flat bottom, tissue culture (TC) grade (Nest, catalog number: 701001)
4. Cell culture dishes,60 mm × 15 mm (NEST, catalog number: 705001)
5. Conical-bottom centrifuge tubes, 50 mL (Uniparts, catalog number: 32117F)
6. Conical-bottom centrifuge tubes, 15 mL (Uniparts, catalog number: 34117F)
7. Microcentrifuge tubes, 1.5 mL (Axygen, catalog number: MCT-150-C)
8. 1,000 μL pipette tips (Uniparts, catalog number: 51131)
9. 200 μL pipette tips (Uniparts, catalog number: 51121Y)
10. 10 μL pipette tips (Axygen, catalog number: T-300)
11. MF-Millipore 0.22 mm MCE membrane, 47 mm (Catalog number: GSWP04700)
12. Nitrile gloves (Kirkland Signature, catalog number: B0CJ4JMTQQ)
13. Serological pipettes 2 mL (Corning, 4486)

### Solutions

1. Culture medium for MDA-MB-231 cells (DMEM complete medium) (see Recipes)
2. 1.5% agarose solution (see Recipes)
3. 1× PBS solution (see Recipes)
4. Trypticase soy broth (TSB) medium (see Recipes)
5. 0.04% Trypsin-EDTA (see Recipes)
6. 500 μM Actinomycin D
7. 100 μg/mL Gentamicin
8. 1 mg/mL Resazurin stock solution
9. 0.15 mg/mL Resazurin stock solution

## Recipies

### a) Culture medium for MDA-MB-231 cells Dulbecco’s modified Eagle medium (DMEM-complete medium)

a) Culture medium for MDA-MB-231 cells Dulbecco's modified Eagle medium (DMEM-complete medium)
| Reagent | Final concentration | Quantity |
| --- | --- | --- |
| DMEM powder | — | As required |
| Sodium bicarbonate (NaHCO<sub>3</sub>) | 3 g/L | 3 g |
| FBS | 5% (v/v) | 50 mL |
| BCS | 5% (v/v) | 50 mL |
| Penicillin–streptomycin | 1% (v/v) | 10 mL |
| Amphotericin B | 0.05% (v/v) | 500 µL |
| L-glutamine (200 mM) | 0.05% (v/v) | 500 µL |
| Sterile distilled water | — | To 1 L final volume |

1. Dissolve DMEM powder in sterile distilled water and stir until completely dissolved.
2. Add 3 g sodium bicarbonate (NaHCO_3_).
3. Adjust the pH to 7.4 by carefully adding drops of NaOH (1 M) or HCl (1 M), as needed.

Under a laminar flow hood, supplement the medium with the following components:

1. 5% Fetal bovine serum (FBS) (50 mL)
2. 5% Bovine calf serum (BCS) (50 mL)
3. 1% Penicillin–streptomycin (10 mL)
4. 0.05% Amphotericin B (500 µL)
5. 0.05% L-Glutamine 200 mM (500 µL)
6. Sterilize the media by filtration using a 0.22 mm MCE membrane in a vacuum filtration system.

**Note:** MDA-MB-231 cells are routinely maintained in this medium. Once prepared, sterilize the complete medium using a 0.22 µm MCE vacuum filtration unit and refrigerate at 4 °C; use within 4 weeks.

### b) 1.5 % agarose solution

| Reagent | Final concentration | Quantity |
| --- | --- | --- |
| Agarose | 1.5% (w/v) | 1.5 g |
| Triple-distilled water | — | To 100 mL |
| Final volume | — | 100 mL |

1. Add 1.5 g agarose to 100 mL triple-distilled water in a glass flask.
2. Sterilize the solution by autoclaving for 15 minutes.

*Note: This agarose solution is used to generate non-adherent surfaces in standard 96-well plates. The prepared solution should be sterilized by autoclaving and used fresh or maintained at ∼60–70 °C until dispensing*.

### c) Trypticase soy broth (TSB) medium

| Reagent | Final concentration | Quantity |
| --- | --- | --- |
| Trypticase soy broth | 1% (w/v) | 30 g |
| Distilled water | — | Up to 1 L |

1. Dissolve all components in approximately 800 mL of distilled water using a magnetic stirrer.
2. Transfer the solution to a 1 L volumetric flask and adjust the final volume to 1 L with distilled water.
3. Sterilize by autoclaving at 120 °C for 20–30 minutes.
4. Allow the solution to cool to room temperature before use. Store appropriately if not used immediately.

### d) 1× PBS solution, pH 7.4

| Reagent | Final concentration | Quantity |
| --- | --- | --- |
| Sodium chloride (NaCl) | 137 mM | 8.0 g |
| Potassium chloride (KCl) | 2.7 mM | 0.2 g |
| Disodium phosphate (Na<sub>2</sub>HPO<sub>4</sub>) | 10 mM | 1.44 g |
| Monopotassium phosphate (KH<sub>2</sub>PO<sub>4</sub>) | 1.8 mM | 0.24 g |
| Triple-distilled water | — | To 1 L |
| Final volume | — | 1 L |

1. Dissolve all components in approximately 800 mL of distilled water using a magnetic stirrer until the solution is clear and no particulates remain.
2. Calibrate the pH meter using standard buffer solutions, then titrate the solution to pH 7.4 by adding 1 M HCl or 1 M NaOH dropwise with continuous stirring. Allow the reading to stabilize after each addition.
3. Transfer the solution to a 1 L volumetric flask and adjust the final volume to 1 L with distilled water. Mix thoroughly by inverting the flask several times.
4. Transfer the solution to an autoclave-safe bottle, loosen the cap slightly to allow pressure equilibration, and autoclave at 121 °C for 20–30 minutes.
5. After autoclaving, allow the solution to cool to room temperature before use. Store appropriately if not used immediately, and label with the solution name, pH and preparation date. Note: Alternatively, a commercial PBS solution may be used.

### e) 0.04 % Trypsin-EDTA

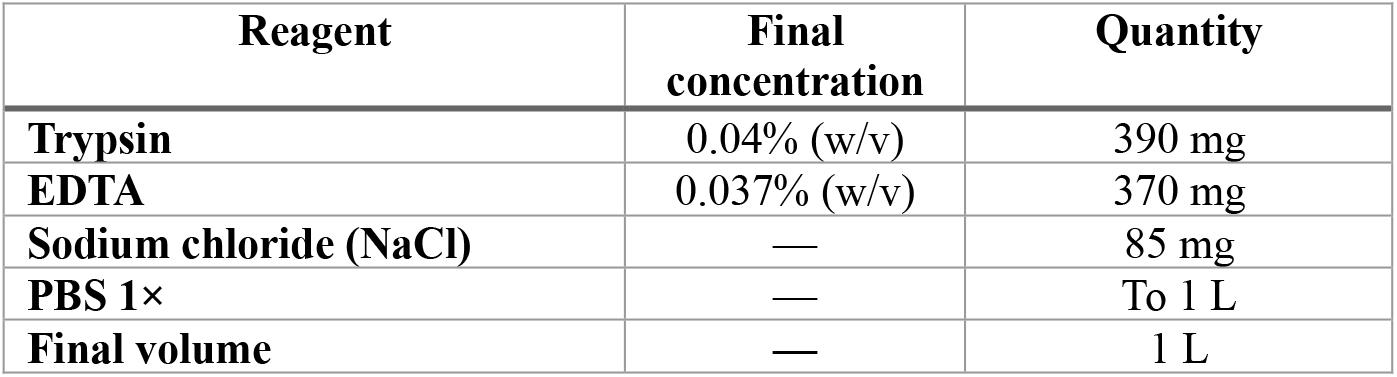

| Reagent | Final concentration | Quantity |
| --- | --- | --- |
| Trypsin | 0.04% (w/v) | 390 mg |
| EDTA | 0.037% (w/v) | 370 mg |
| Sodium chloride (NaCl) | — | 85 mg |
| PBS 1× | — | To 1 L |
| Final volume | — | 1 L |

1. Prepare 1 L of sterile 1× phosphate-buffered saline (PBS) according to standard procedure.
2. In a separate sterile container, dissolve 390 mg trypsin and 85 mg NaCl in 10 mL of the prepared 1× PBS. Mix gently until completely dissolved.
3. To the remaining PBS (∼990 mL), add 370 mg EDTA and stir until fully dissolved.
4. Combine the trypsin-NaCl solution with the EDTA-containing PBS and stir thoroughly to ensure complete mixing.
5. Under aseptic conditions, filter-sterilize the combined solution using a 0.22 µm vacuum filtration unit.
6. Aliquot the sterile solution into appropriate volumes, label clearly, and store at –20 °C until use.

*Note: This trypsin–EDTA solution is used for cell detachment. The prepared solution is sterilized by filtration (0*.*22 µm) and stored at −20 °C until use*.

### e) 500 µM actinomycin D stock solution

**f) 500 $\mu\text{M}$ actinomycin D stock solution**
| Reagent | Final concentration | Quantity |
| --- | --- | --- |
| Actinomycin D | 500 $\mu\text{M}$ | As required |
| DMSO | — | As required |

Safety precautions: Actinomycin D is a potent cytotoxic and teratogenic agent. Wear appropriate personal protective equipment (PPE), including gloves and safety goggles, and handle all materials in a designated chemical fume hood or biological safety cabinet.

1. Working under aseptic conditions in a cell culture flow hood, dissolve the appropriate amount of Actinomycin D in sterile DMSO to achieve a final concentration of 500 µM.
2. Mix gently by pipetting or vortexing until completely dissolved.
3. Dispense the solution into amber or foil-wrapped microcentrifuge tubes to protect from light.
4. Label each aliquot clearly with the compound name, concentration and date of preparation.
5. Store protected from light at −20 °C until use. Avoid repeated freeze-thaw cycles.

### g) Gentamicin stock solution (100 µg/mL)

**g) Gentamicin stock solution (100 $\mu\text{g}/\text{mL}$ )**
| Reagent | Final concentration | Quantity |
| --- | --- | --- |
| Gentamicin sulfate | 100 $\mu\text{g}/\text{mL}$ | As required |
| Sterile distilled water | — | As required |

Common working stock are 10 mg/mL or 50 mg/mL. Adjust the volume of water accordingly based on the mass of gentamicin powder used.

1. Dissolve the required mass of gentamicin sulfate powder in an appropriate volume of sterile distilled water to achieve the desired stock concentration (e.g., 10 mg/mL or 50 mg/mL). Mix gently until completely dissolved.
2. Under aseptic conditions in a cell culture flow hood, sterilize the solution by filtration through a 0.22 µm syringe filter.
3. Aliquot the sterile solution into appropriately sized, sterile microcentrifuge tubes or vials.
4. Label each aliquot with the antibiotic name, concentration and date of preparation.
5. Store at 4 °C for short-term use (up to one month) or at −20 °C for long-term storage. Avoid repeated freeze-thaw cycles.

### h) Resazurin stock solution (0.15% w/v)

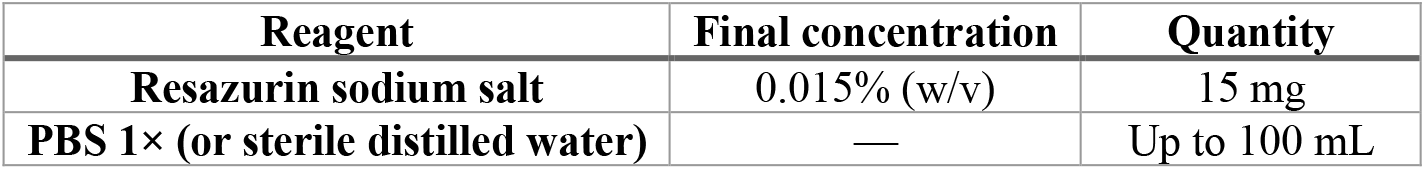

### Resazurin stock solution (1 mg/mL)

***Note:*** This protocol yields a 1 mg/mL resazurin stock solution, commonly used as a cell viability indicator in proliferation and cytotoxicity assays. This concentration is typically used as a 10× or 100× stock depending on the assay format.

1. Accurately weigh the required mass of resazurin sodium salt using an analytical balance. For a 1 mg/mL stock, weigh 100 mg of resazurin sodium salt to prepare 100 mL of solution.
2. Transfer the powder to a sterile glass beaker or flask and dissolve in approximately 80 mL of sterile phosphate-buffered saline (PBS).
3. Stir gently on a magnetic stirrer or swirl by hand until the powder is completely dissolved, yielding a uniform blue solution.
4. Transfer the solution to a 100 mL volumetric flask and adjust the final volume to 100 mL with sterile PBS. Mix thoroughly by inverting several times.
5. Immediately cover the container with aluminum foil or use an amber bottle to protect the solution from light, as resazurin is photosensitive.
6. Under aseptic conditions in a cell culture flow hood, filter-sterilize the solution through a 0.22 µm membrane filtration unit.
7. Aliquot the sterile solution into light-protected tubes (amber microcentrifuge tubes or clear tubes wrapped in aluminum foil).
8. Label each aliquot with the solution name, concentration (1 mg/mL) and date of preparation.
9. Store protected from light at −20 °C until use. Avoid repeated freeze-thaw cycles.

### Preparation of resazurin working solution (0.15 mg/mL) from 1 mg/mL stock

***Note:*** This protocol yields a 0.15 mg/mL (150 µg/mL) resazurin working solution, typically used as an intermediate dilution for cell viability assays. This concentration is often used as a 10× stock, diluted 2:10 in cell culture medium to achieve a working concentration of 30 µg/mL.

1. Calculate the required dilution factor using the formula: C_1_V_1_ = C_2_V_2_, where:
  - C_1_ = Initial concentration (1 mg/mL)
  - C_2_ = Desired concentration (0.15 mg/mL)
  - V_2_ = Desired final volume of working solution
  - V_1_ = Volume of stock solution needed

### Example calculation for 10 mL of working solution

○ (1 mg/mL) × V_1_ = (0.15 mg/mL) × (10 mL)
○ V_1_ = (0.15 × 10) / 1 = 1.5 mL of stock solution

2. Under aseptic conditions in a cell culture flow hood, aliquot the calculated volume of 1 mg/mL resazurin stock solution into a sterile container.
3. Add sterile 1× PBS (or sterile cell culture medium, depending on your application) to achieve the desired final volume. For 10 mL: Add 1.5 mL of resazurin stock (1 mg/mL) + 8.5 mL of sterile 1× PBS.
4. Mix gently by pipetting or swirling until thoroughly combined.
5. Protect the solution from light by using an amber tube or covering with aluminum foil.
6. If not used immediately, aliquot into light-protected tubes and label with the concentration (0.15 mg/mL) and date.
7. Store protected from light at 4 °C for short-term use (up to 2 weeks) or at −20 °C for longer storage. Avoid repeated freeze-thaw cycles.

## Equipment

1. Centrifuge (PowerSpin™, model: C856)
2. Micropipettes (Axygen, AP-10, AP-100 and AP-1000):
  a. 0.5–10 µL Single-channel Pipettor (Axygen® Axypet®, catalog number: AP-10)
  b. 10–100 µL Single-channel Pipettor (Axygen® Axypet®, catalog number: AP-100)
  c. 100–1,000 µL Single-channel Pipettor (Axygen® Axypet®, catalog number: AP-1000)
3. pH meter (Apera, PH700)
4. Orbital shaker (Benchmark, model: BT302)
5. Digital stirring hot plate (Thermo Scientific, model: SP131015Q)
6. Inverted microscope (Carl Zeiss, model: 37081)
7. Laminar flow cabinet (Thermo Fisher Scientific, model: 1340)
8. Incubator for cell culture (Thermo Fisher Scientific, model: 3422)
9. Neubauer chamber (Mariendfeld, catalog number: 0610010)
10. Microwave oven (standard laboratory or domestic units, 600-100 W) for agarose melting.
11. Vacuum filtration system (Nalgene, catalog number 300-4050)
12. Incubator for bacterial culture (Thermo Fisher Scientific, model:SHKE6000)
13. Microplate reader Varioskan Flash (Thermo Scientific, model: N06354)

### Software and datasets

1. ImageJ (https://imagej.net/ij/download.html)
2. GraphPad Prism 9.1.1 (https://www.graphpad.com/features)

### Procedure

Prior to initiating any experimental procedures, prepare the biosafety cabinet as follows:

1. Thoroughly disinfect all work surfaces by wiping with 70% ethanol.
2. Turn on the ultraviolet (UV) lamp and expose the cabinet for 15 minutes to ensure surface sterilization.
3. After UV exposure, turn off the UV lamp and activate the laminar airflow. Allow the cabinet to run for 2–3 minutes before introducing materials.

All cell culture and bacterial manipulations must be performed under strict aseptic conditions within the biosafety cabinet, using only sterile materials and proper sterile technique.

#### A. Assessment of bacterial growth inhibition by gentamicin using a resazurin-based viability assay (Fig. 1)

**Figure. 1.**
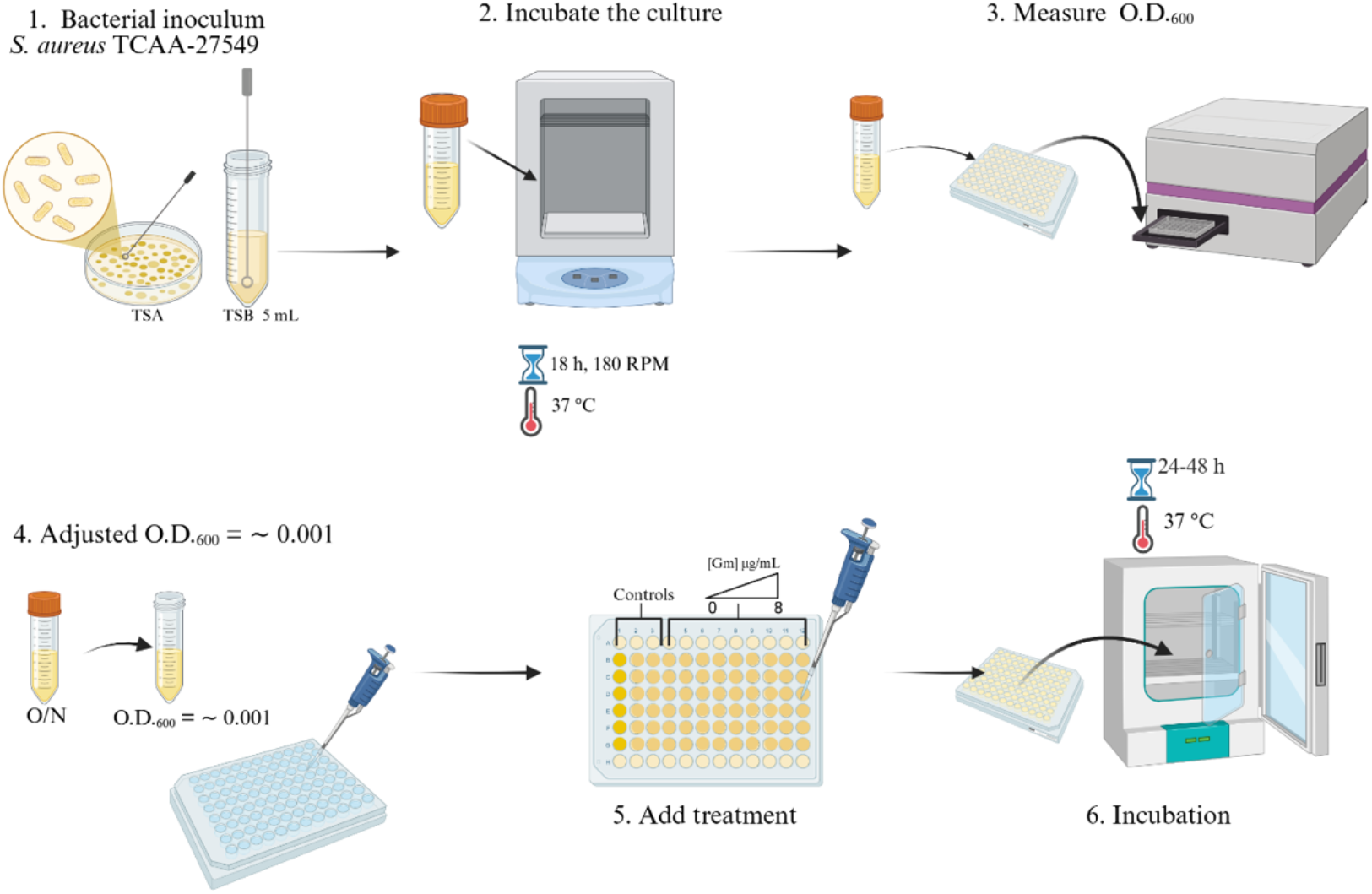
Bacterial culture optimization and subsequent gentamicin treatment for viability assessment.

##### I. Bacterial culture and inoculum preparation

1. Using a sterile loop, inoculate 5 mL of tryptic soy broth (TSB) supplemented with 1% glucose with a single colony of *Staphylococcus aureus* TCAA-27549 obtained from an agar plate.
2. Incubate the culture at 37 °C with shaking at 180 rpm for 18 hours.
3. Measure the optical density (OD) at 600 nm using a Varioskan Flash microplate reader.
4. Following incubation, adjust the bacterial suspension to a final concentration of 5 × 10^5^ CFU/mL (O.D. ^600^ = ∼0.001) using fresh TSB.
5. Dispense 100 µL of the adjusted bacterial suspension into each well of a sterile 96-well microplate.

4. Prepare a series of gentamicin working solutions (0–8 µg/mL) by diluting a 100 µg/mL stock solution in sterile water.
5. Add the appropriate volume of each gentamicin working solution to the designated wells to achieve the desired final concentrations.
6. Include the following controls in each plate:
  - Medium control: Sterile TSB only.
  - No-treatment control: Bacterial suspension without antibiotic.
  - Vehicle: Sterile TSB with gentamicin (to verify sterility).
7. Incubate the plates at 37 °C for 24 or 48 hours, depending on the experimental requirements.

##### II. Viability assessment using resazurin (Fig. 2)

**Figure 2.**
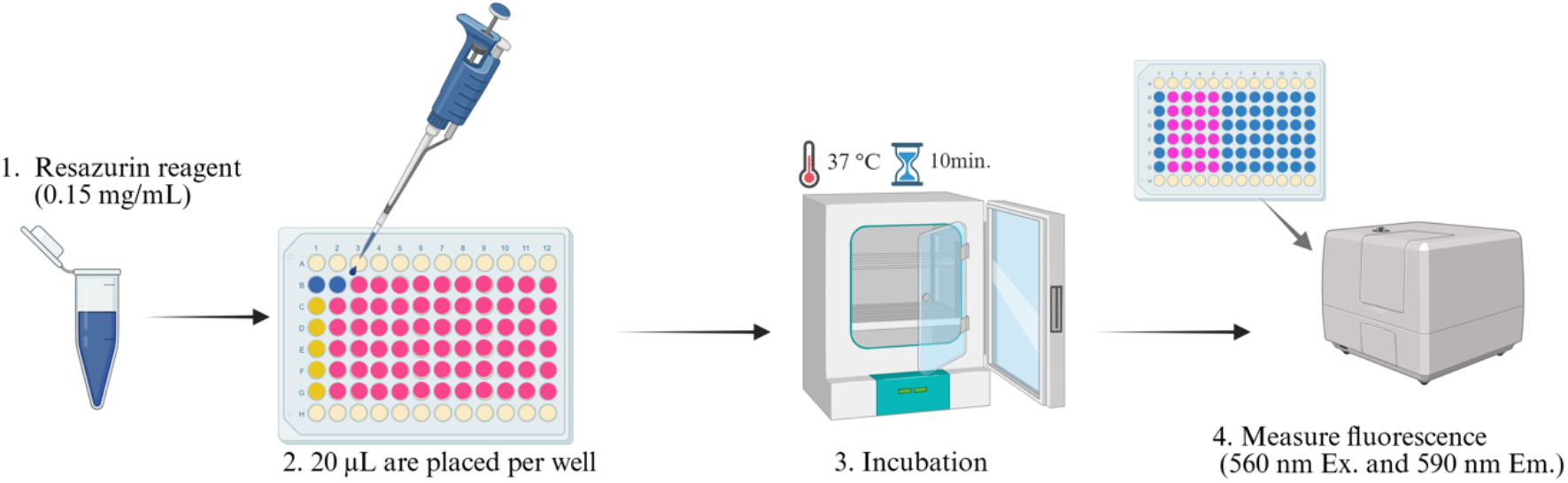
Resazurin reduction assay for bacterial growth inhibition.

1. After the incubation period, add 20 µL of resazurin working solution (0.15 mg/mL) to each well.
2. Incubate the plates at 37 °C for 10 minutes, protected from light, to allow resazurin to be reduced to resorufin by viable baccteria.
3. Measure fluorescence using a microplate reader with excitation and emission wavelengths set to 560 nm and 590 nm, respectively. Record the fluorescence intensity as an indicator of bacterial viability (Fig. 2 and 3).

**Figure 3.**
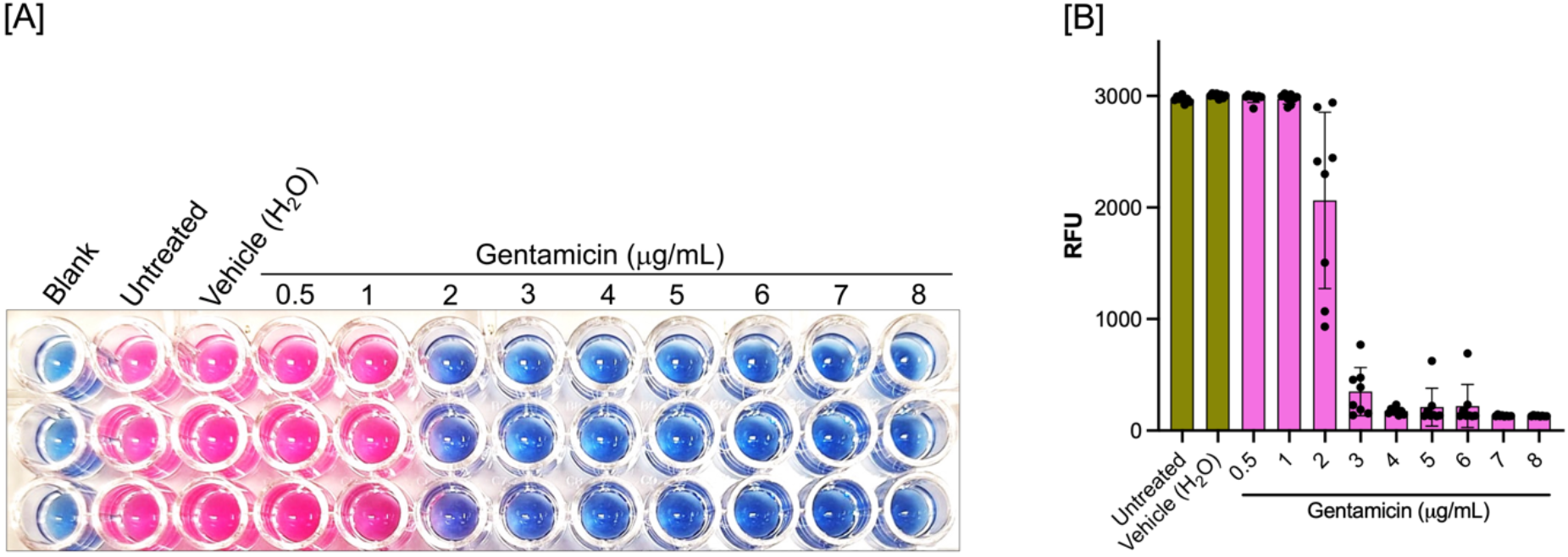
Bacterial viability (*S. aureus* ATCC-27549). (A) Colorimetric reaction of bacterial cultures after 24 hours of exposure to different concentrations of gentamicin (0.5, 1, 2, 3, 4, 5, 6, 7, and 8 µg/mL). (B) Quantification of bacterial viability through fluorescence measurements, expressed as relative fluorescence units (RFU). Data are presented as mean ± standard deviation (SD); n = 8 biological replicates.

#### B. 2D Cell culture protocol for MDA-MB-231 cells (Fig. 4)

**Figure. 4.**
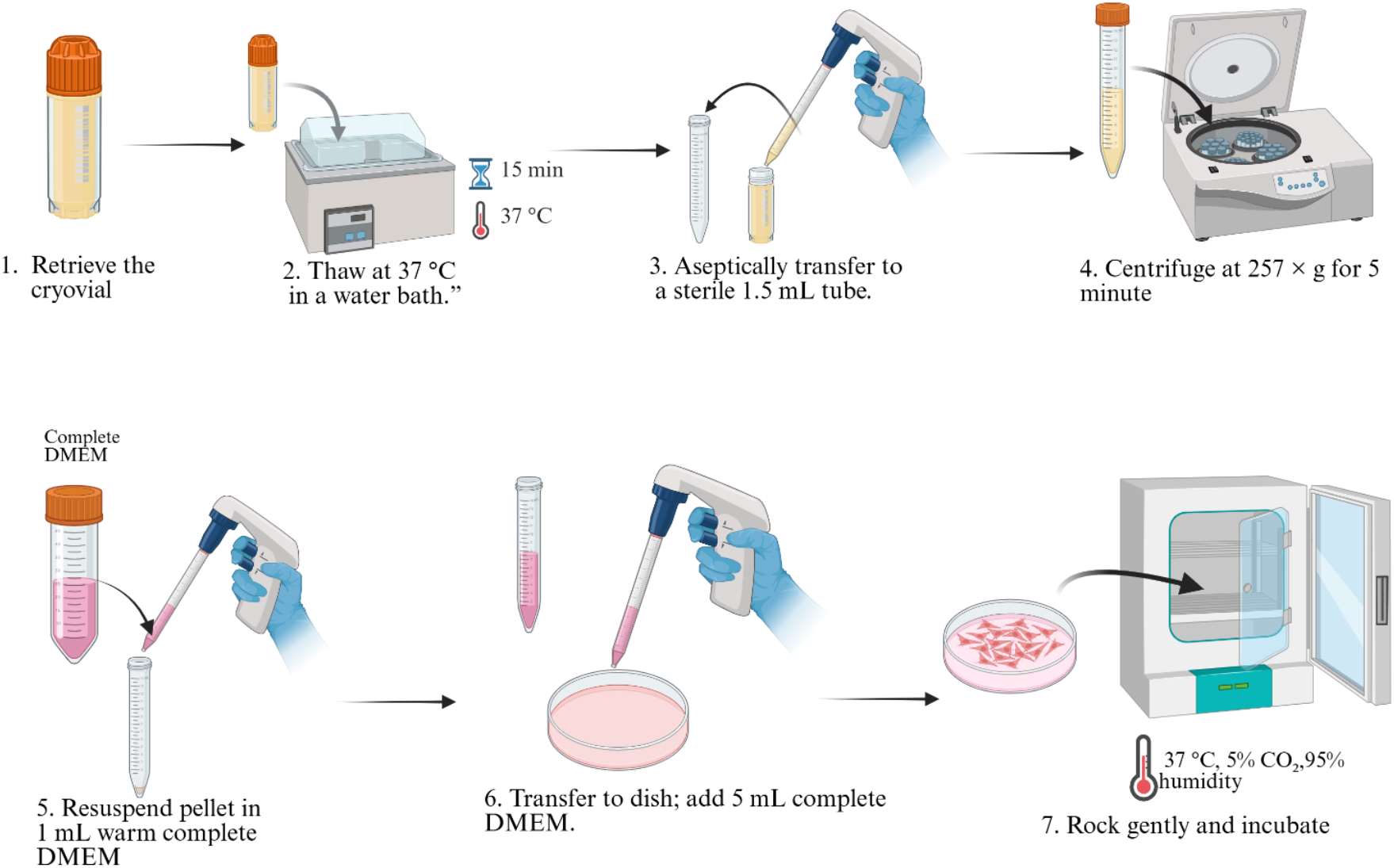
MDA-MB-231 cells cultured in 2D monolayer.

##### I. Thawing MDA-MB-231 cells

1. Retrieve the cryovial containing MDA-MB-231 cells from liquid nitrogen storage.
2. Immediately place the vial in a 37 °C water bath and gently swirl until the cell suspension is completely thawed (approximately 1–2 minutes).
3. Under aseptic conditions in a biosafety cabinet, transfer the thawed cell suspension to a sterile 1.5 mL tube using a sterile 2 mL serological pipette.
4. Centrifuge at 257 × g for 5 minutes at room temperature to pellet the cells.
5. Carefully aspirate the supernatant using a sterile 2 mL serological pipette, avoiding disturbance of the cell pellet.
6. Gently resuspend the cell pellet in 1 mL of fresh, pre-warmed complete DMEM medium by pipetting up and down.
7. Transfer the cell suspension to a 60 mm × 15 mm cell culture dish and add an additional 5 mL of complete DMEM medium.
8. Rock the dish gently in a crosswise motion to distribute cells evenly, then place in a humidified incubator at 37 °C with 5% CO_2_.

**Note:** RPMI medium may be used as an alternative to DMEM, depending on experimental requirements.

##### II. Subculture and preparation of MDA-MB-231 cells

1. Culture MDA-MB-231 cells in 60 mm × 15 mm cell culture dishes containing complete DMEM medium at 37 °C with 5% CO_2_ until they reach approximately 80% confluence.
2. Aspirate the spent culture medium carefully using a sterile serological pipette.
3. Wash the cell monolayer gently with 5 mL of sterile 1× PBS to remove any residual serum that may inhibit trypsin. Aspirate and discard the PBS.
4. Add 3 mL of pre-warmed 0.25% trypsin-EDTA solution to completely cover the cell monolayer completely.
5. Incubate the dish at 37 °C for 3–5 minutes. Observe periodically under an inverted microscope until cells have rounded up and detached from the surface.
6. Neutralize the trypsin by adding 6 mL of complete DMEM medium (containing serum) to the dish.
7. Gently pipette the cell suspension up and down to break up any clumps, then transfer the entire volume to a sterile 15 mL conical tube.
8. Centrifuge at 257 × g for 5 minutes at room temperature.
9. Carefully aspirate the supernatant without disturbing the cell pellet.
10. Resuspend the cell pellet in 1 mL of fresh complete DMEM medium by gentle pipetting.
11. Perform a cell count before seeding for experiments or further passage.

**Note 1:** RPMI medium may be used as an alternative to DMEM.

**Note 2:** Cells should be passaged at least three times after thawing before use in experiments to ensure recovery and stable growth.

##### III. Cell counting with a Neubauer chamber (Fig. 5)

**Figure. 5.**
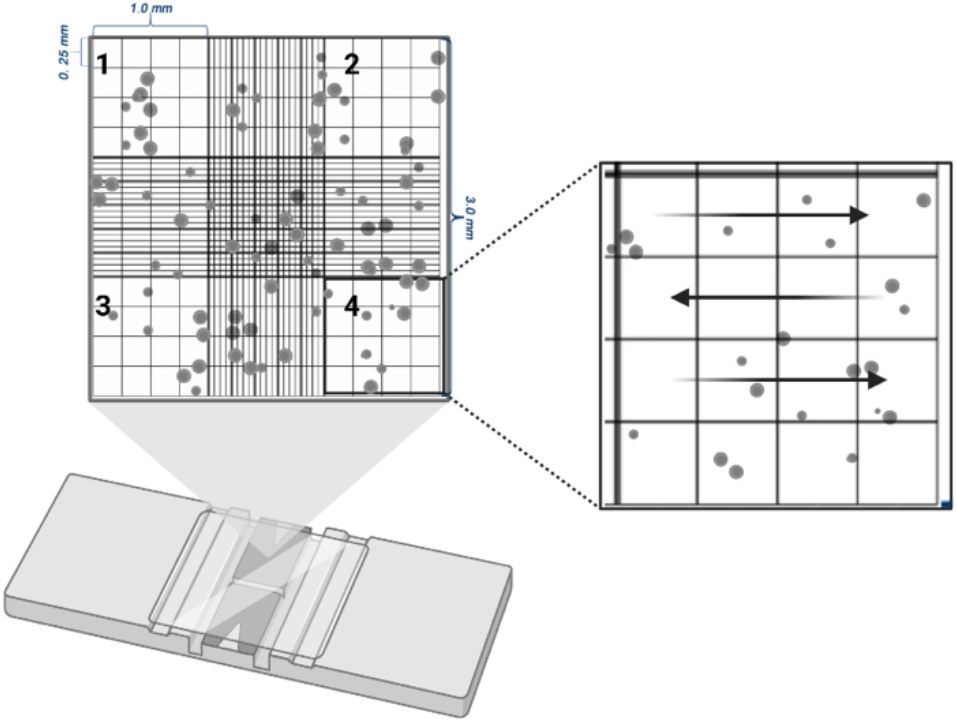
Neubauer chamber cell counting. (Left) Four corner quadrants used for counting. (Right) Counting rule: include cells touching top/left borders, exclude those touching bottom/right.

1. Prepare a 1:10 dilution of the cell suspension for counting:
  - Add 90 µL of complete DMEM medium to a sterile 1.5 mL microcentrifuge tube.
  - Add 10 µL of the well-mixed cell suspension to the tube.
  - Mix gently by pipetting.
2. Pipette 10 µL of the diluted cell suspension into the chamber of a Neubauer hemocytometer, allowing the liquid to be drawn in by capillary action. Ensure the chamber is fully covered without overflow.
3. Using an inverted microscope, count the cells in the four corner squares (each consisting of 16 small squares) of the hemocytometer grid. Include cells touching the upper and left boundaries; exclude those touching the lower and right boundaries.
4. Calculate the cell concentration using the following formula:

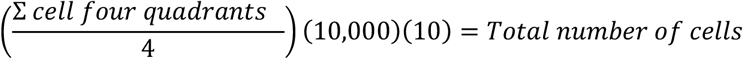

Example calculation:

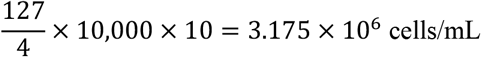

##### IV. Cell synchronization

After allowing cells to attach and grow for 24 hours post-seeding:

1. Carefully aspirate the culture medium from each well using a sterile pipette tip connected to a vacuum line, taking care not to disturb the cell monolayer.
2. Gently wash each well once with 100 µL of PBS 1× to remove residual serum. Aspirate and discard the wash.
3. Add 100 µL of fresh incomplete DMEM to each well.
4. Incubate the plate at 37 °C in a humidified incubator with 5% CO_2_ for 24–48 hours to arrest cells in a quiescent state prior to treatment.

**Note:** The duration of synchronization may need optimization depending on cell line and experimental conditions.

##### V. Treatment with actinomycin D

1. After the synchronization period, carefully aspirate the incomplete DMEM from each well.
2. Add 100 µL of Actinomycin D working solutions prepared in incomplete DMEM (or appropriate medium) to achieve final concentrations ranging from 2 to 14 µM.
3. Include the following control wells in each experiment:
  - No-treatment control: Cells with complete medium only (no drug).
  - Vehicle control: Cells with medium containing the same concentration of solvent (e.g., DMSO) used to dissolve Actinomycin D.
4. Incubate the plate at 37 °C with 5% CO_2_ for 24–48 hours, depending on the experimental time course.

##### VI. Cell viability assessment using resazurin (Fig. 6)

**Figure 6.**
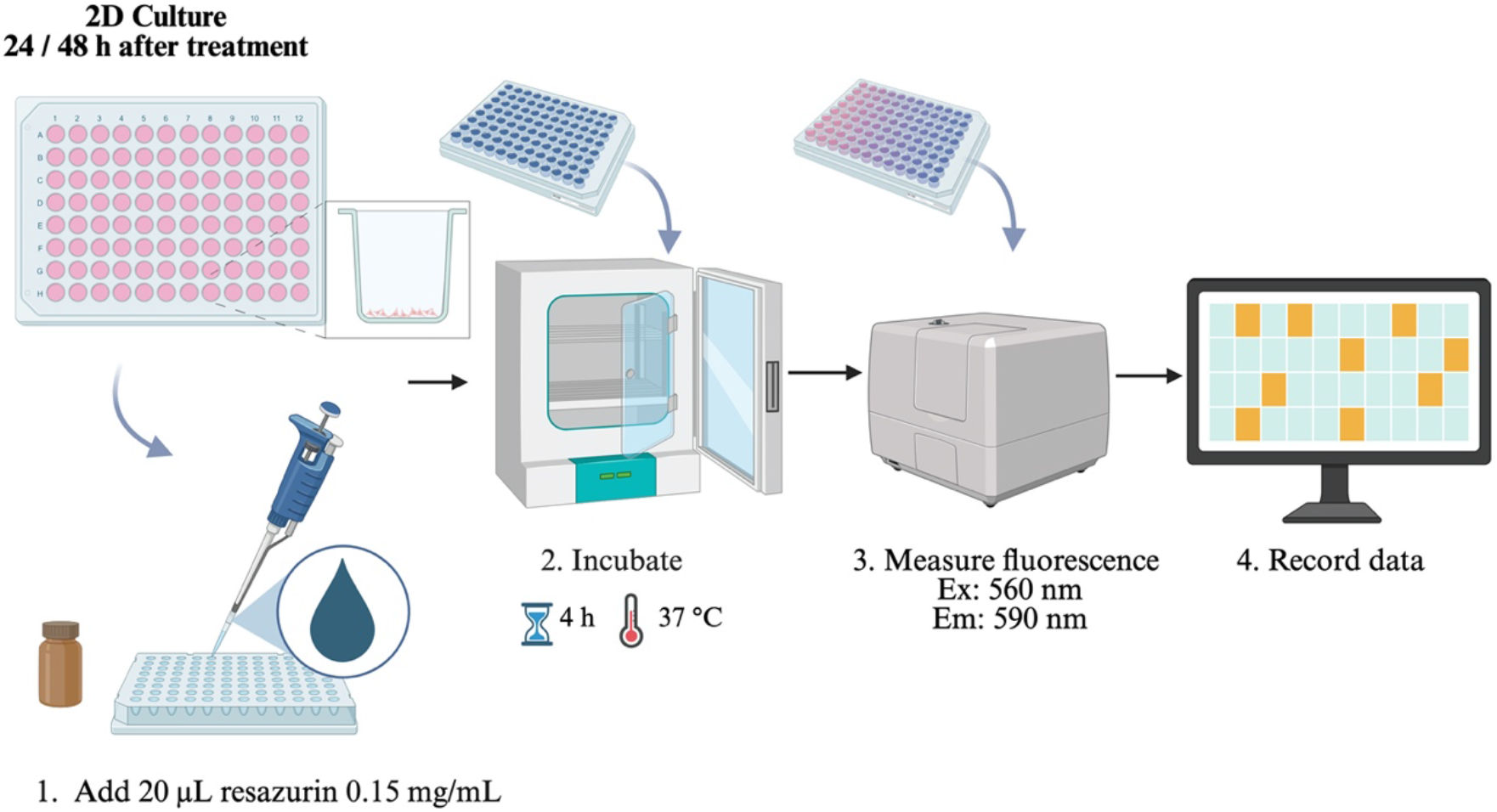
Resazurin reduction assay for eukaryotic cell viability in 2D culture.

1. After the treatment incubation period, add 20 µL of resazurin working solution (0.15 mg/mL in sterile PBS 1×) directly to each well.
2. Gently tap the plate to mix, taking care not to introduce bubbles.
3. Incubate the plate at 37 °C for 2–4 hours, protected from light, to allow viable cells to reduce resazurin (blue, non-fluorescent) to resorufin (pink, highly fluorescent).
4. Measure fluorescence intensity using a microplate reader with excitation and emission wavelengths set to 560 nm and 590 nm, respectively.
5. Record fluorescence values as an indicator of cell viability. Higher fluorescence corresponds to greater metabolic activity and thus higher viability (Fig. 7).

**Figure 7.**
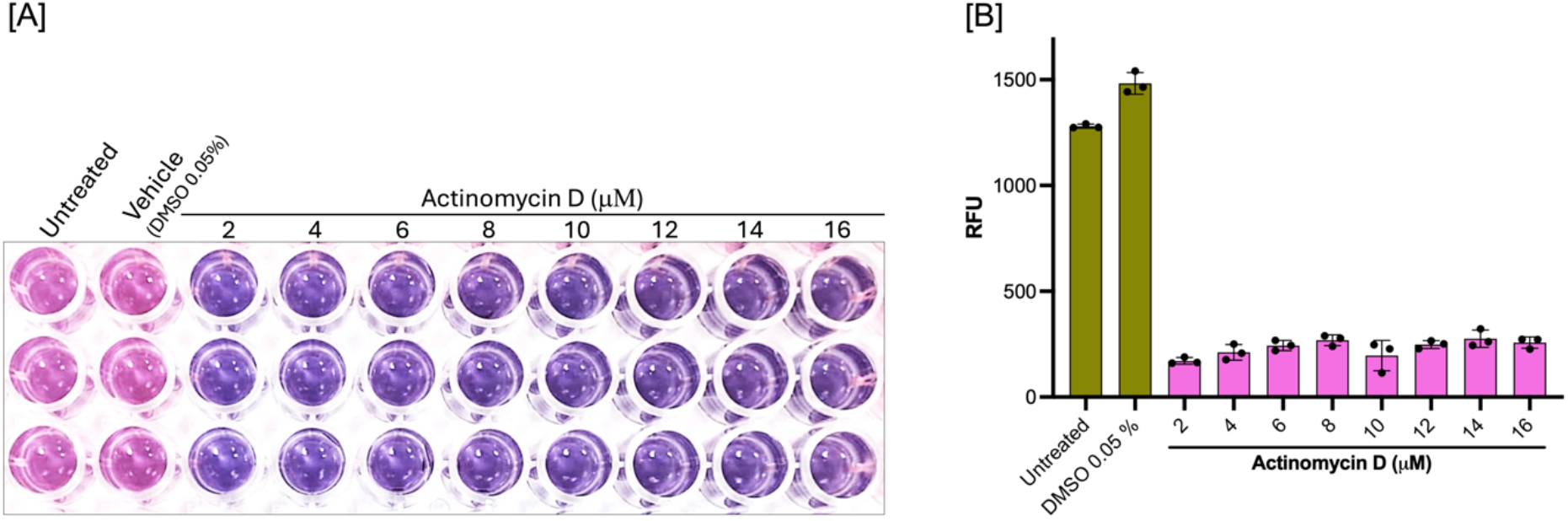
Viability of eukaryotic cells (MDA-MB-231) in 2D culture. (A) Colorimetric assay with resazurin of MDA-MB-231 cells after 24 hours of exposure to increasing concentrations of actinomycin D (2, 4, 6, 8, 10, 12, 14, and 16 µM). (B) Quantification of cell viability based on fluorescence measurements, expressed as relative fluorescence units (RFU). Data are presented as mean ± standard deviation (SD); n = 3 biological replicates.

**Note:** The optimal incubation time for resazurin may vary; monitor color change visually and adjust incubation time if necessary (typically 1–4 hours).

#### C. 3D cell culture protocol for MDA-MB-231 spheroid formation and treatment (Fig. 8)

**Figure. 8.**
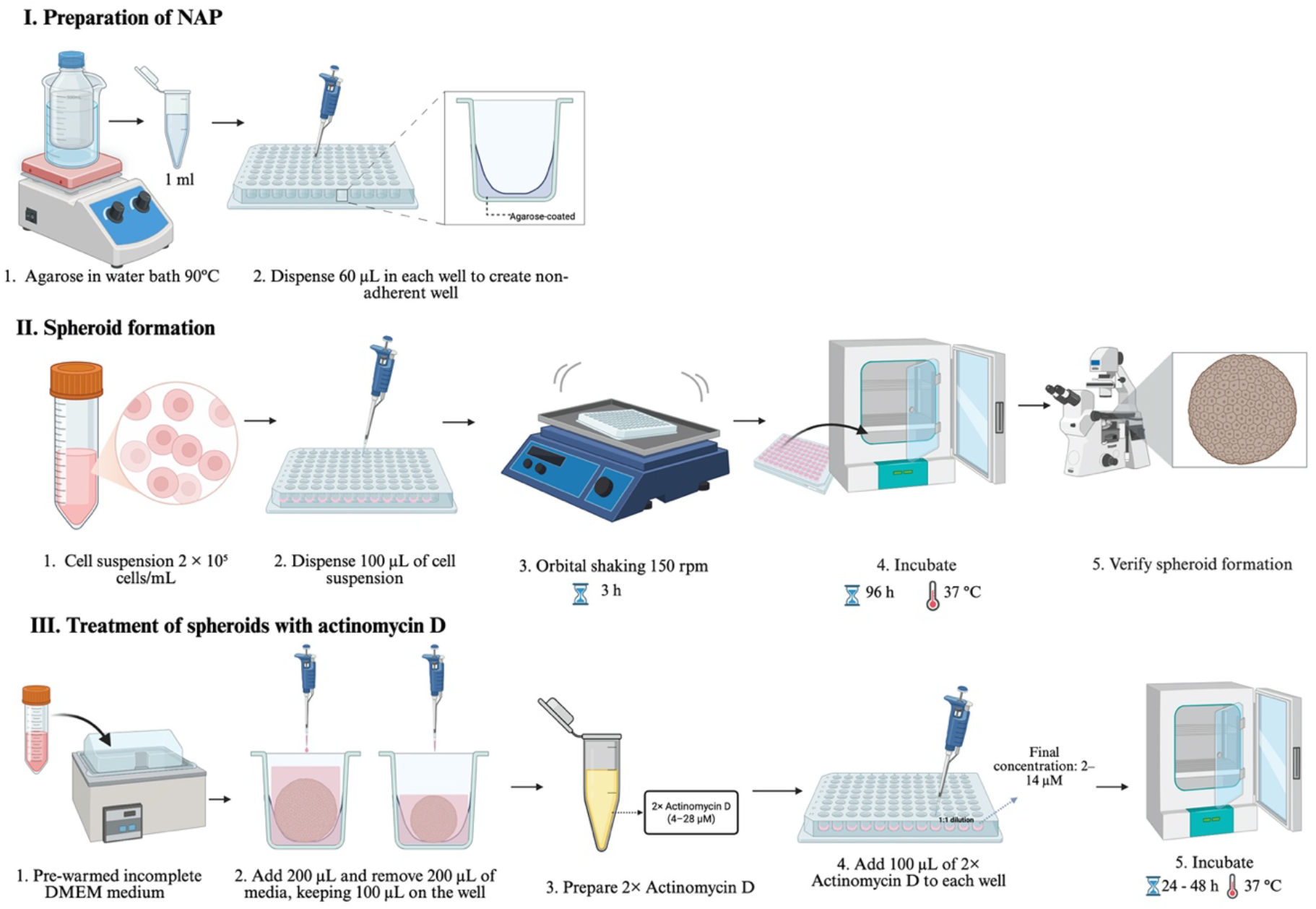
Workflow for MDA-MB-231 spheroid generation. Schematic representation of the steps involved in obtaining 3D spheroids from MDA-MB-231 breast cancer cells.

##### I. Preparation of cost-effective non-adherent plates using agarose coating

1. Prepare a sterile 1.5% (w/v) agarose solution according to standard procedures (see recipes). Maintain the molten agarose in a water bath set to 100 °C on a stirring hot plate to prevent solidification.
2. Use a sterile, flat-bottom 96-well cell culture plate for coating.
3. Using strict aseptic technique, transfer 1 mL of the molten 1.5% agarose solution into a sterile 1.5 mL microcentrifuge tube. This serves as a working aliquot to minimize contamination of the main stock.
4. Immediately dispense 60 µL of the molten agarose into each well of the 96-well plate using the working aliquot tube (Fig. 8) (Cervantes-Rivera et al., 2026). **Critical note:** A multichannel pipette is not recommended for this step, as standard plastic reagent reservoirs are not compatible with the high temperature required to keep the agarose molten. Working quickly with a single-channel pipette is essential to prevent agarose solidification in the tip or tube.
5. Repeat Step 4 until all desired wells are filled. If the agarose in the working aliquot begins to solidify, replace it with fresh molten agarose from the main stock.
6. Allow the agarose to solidify undisturbed at room temperature. Once solidified, place the plate uncovered in a cell culture hood for 15 minutes to allow any residual moisture to evaporate under laminar flow.
7. Wrap the plate securely in clean plastic film (e.g., Parafilm) and place it in a sealed plastic bag to prevent dehydration and contamination.
8. Store the prepared agarose-coated plates at 4 °C until use. The plates can be stored for 2–3 weeks under these conditions.

##### II. Spheroid formation

For spheroid formation, follow the standard 2D cell culture procedure (including thawing, subculture, and cell counting using a Neubauer chamber) up to the point of obtaining a single-cell suspension.

1. Prepare a cell suspension in complete DMEM medium at a density of 20,000 cells per 100 µL (equivalent to 2 × 10^5^ cells/mL).
2. Using a micro pipette, gently dispense 100 µL of the cell suspension into each well of the pre-coated 96-well low-attachment plate prepared in Section C-I.
3. To promote cell aggregation and spheroid formation, place the plate on an orbital shaker set to 150 rpm for 3 hours at room temperature or in the incubator.
4. Following the shaking step, incubate the plate statically at 37 °C in a humidified atmosphere containing 5% CO_2_ for 96 hours to allow spheroid maturation.
5. During the incubation period, replace the culture medium every 48 hours. To do so:
  - Carefully aspirate 100 µL of spent medium from each well, acoiding the forming spheroid at the bottom.
  - Gently add 100 µL of fresh, pre-warmed complete DMEM along the wall of the well.
6. After 96 hours, verify successful spheroid formation by observing the wells under an inverted microscope. Spheroids should appear as compact, three-dimensional structures (Figure 8).

Note: The optimal seeding density and incubation time may vary depending on the cell line. Optimization experiments may be required.

##### III. Treatment of spheroids with actinomycin D

1. After spheroid formation is confirmed, carefully add 200 µL of pre-warmed incomplete DMEM to each well by dispensing the liquid slowly along the well wall to avoid displacing the spheroid.
2. Remove 200 µL from each well, keep 100 µL of medium in each well at all time.
3. Prepare 2× concentrated working solutions of Actinomycin D in incomplete DMEM to achieve final concentrations ranging from 2 to 14 µM after addition to the wells. For example, to obtain a final concentration of 10 µM, prepare a 20 µM working solution.

1. Gently add 100 µL of the appropriate 2× Actinomycin D working solution to each well along the wall of the well. This will bring the total volume to 200 µL and achieve the desired final drug concentration (Figure 8).
2. Include the following control wells in each experiment:
  - Untreated control: Spheroids with incomplete DMEM medium only (no drug).
  - Vehicle control: Spheroids with incomplete DMEM medium containing the same concentration of solvent (e.g., DMSO 0.05%) used to dissolve Actinomycin D.
3. Incubate the plate at 37 °C with 5% CO_2_ for 24–48 hours, depending on the experimental time course.

**Note:** The volume of medium removed and added should be carefully controlled to maintain consistent final concentrations. If 100 µL was removed, adding 100 µL of 2× solution is appropriate. If a different volume was removed, adjust accordingly.

##### IV. Cell viability assessment in spheroids using resazurin treatment (Fig. 9)

**Figure 9.**
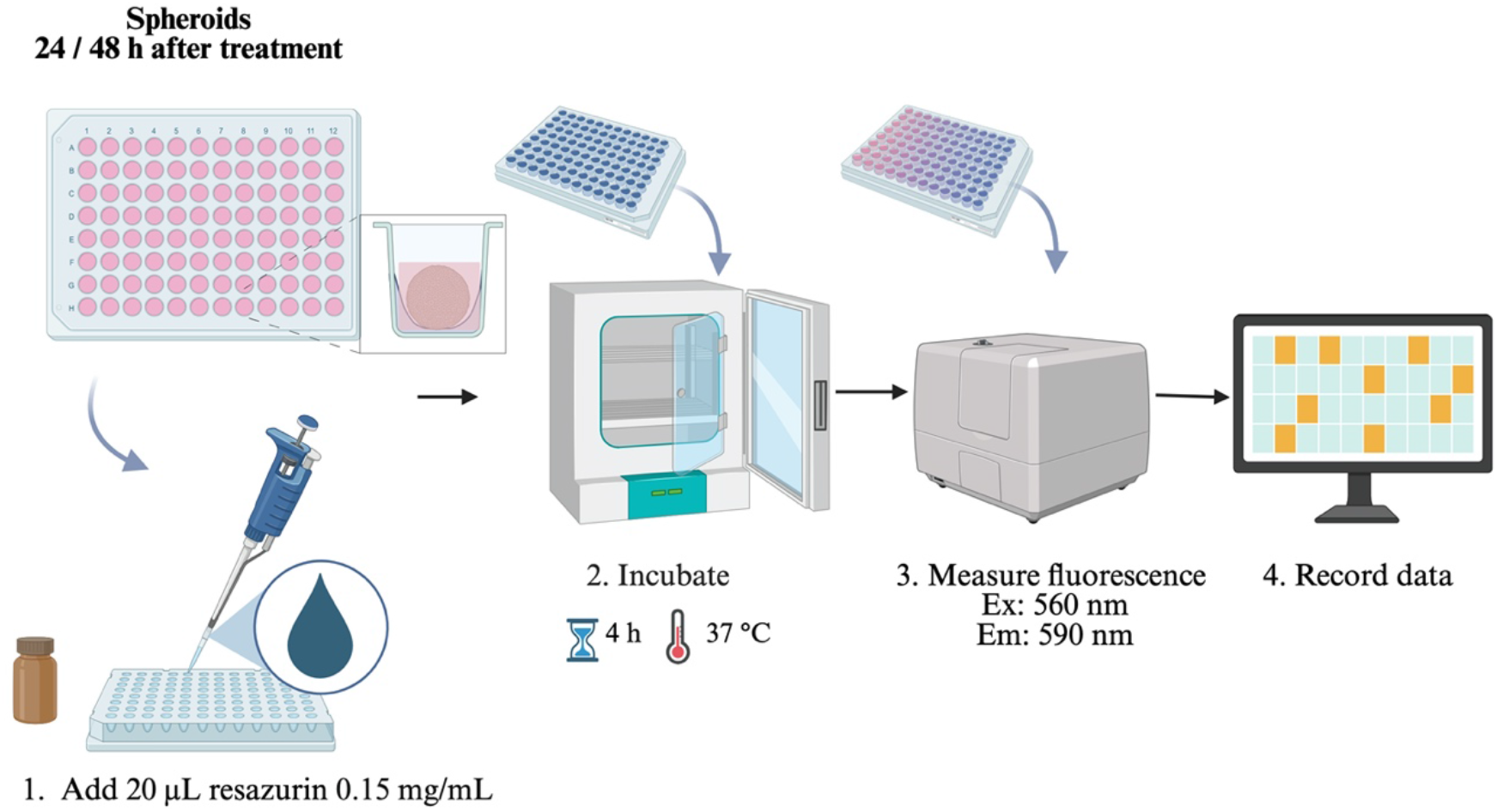
Resazurin reduction assay for eukaryotic cell viability in 3D culture (spheroids).

1. After the treatment incubation period, add 40 µL of resazurin working solution (0.015% w/v in sterile PBS 1×) directly to each well. Critical note: To avoid disturbing or displacing the spheroid, dispense the resazurin solution gently along the wall of the well, not directly onto the spheroid.
2. Gently tap the plate to mix, taking care not to introduce bubbles or disrupt the spheroids.
3. Incubate the plate at 37 °C for 4 hours, protected from light, to allow viable cells within the spheroid to reduce resazurin (blue, non-fluorescent) to resorufin (pink, highly fluorescent).
4. Measure fluorescence intensity using a microplate reader with excitation and emission wavelengths set to 560 nm and 590 nm, respectively.
5. Record fluorescence values as an indicator of spheroid viability. Higher fluorescence corresponds to greater metabolic activity and thus higher viability (Fig. 10).

**Figure 10.**
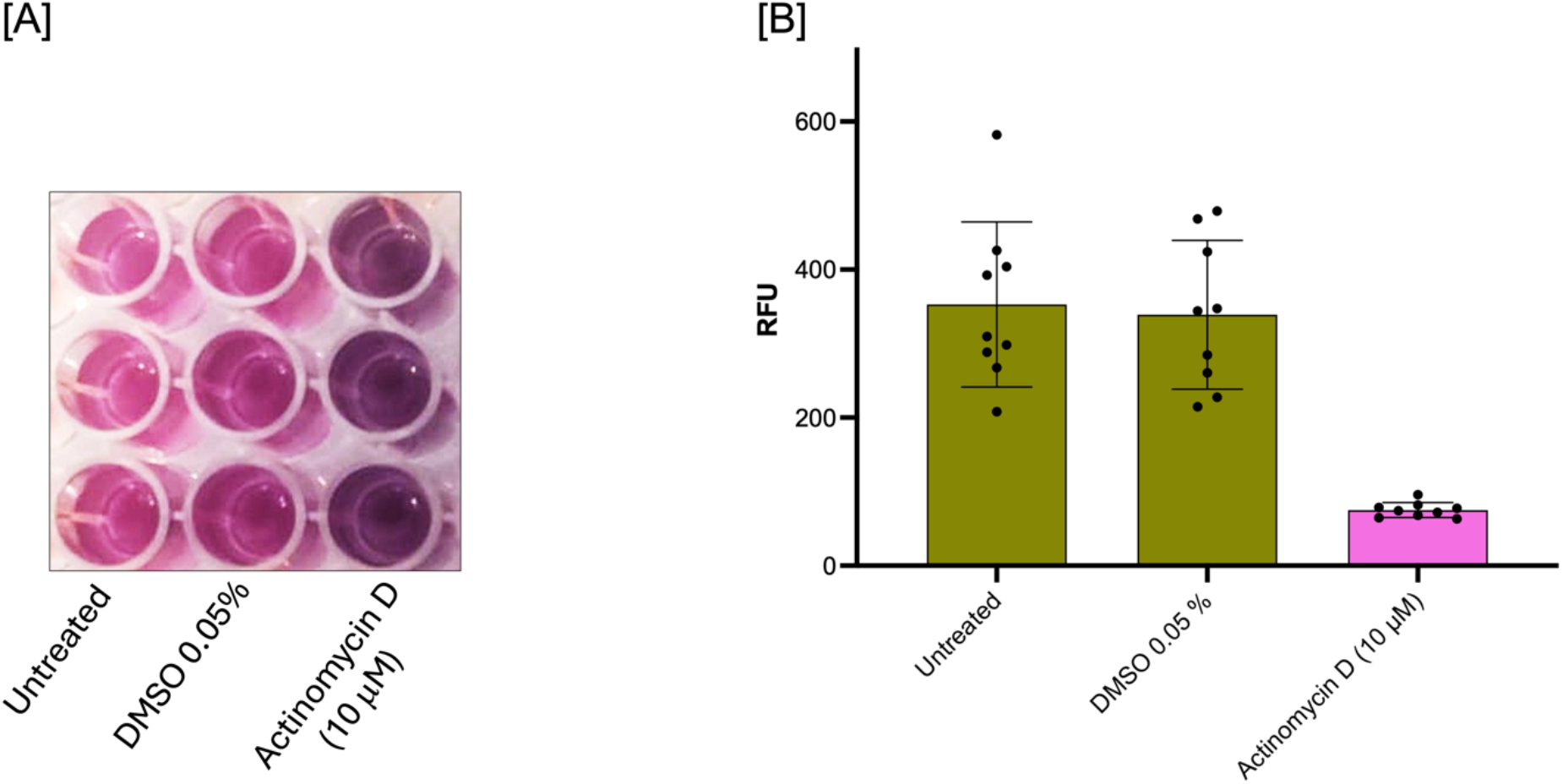
Viability of MDA-MB-231 cells in 3D culture (Spheroids). Spheroids were exposed to actinomycin D (10 µM) for 24 hours. (A) Colorimetric assay with resazurin of cell viability. (B) Quantification of cell viability based on fluorescence measurements, expressed as relative fluorescence units (RFU). Data are presented as mean ± standard deviation (SD); n = 9 biological replicates.

**Note:** The optimal incubation time for resazurin in 3D spheroids may be longer than for 2D cultures due to limited diffusion. A time-course experiment (e.g., 2, 4 and 6 hours) may be necessary to determine the optimal incubation period for your specific spheroid model.

## Validation of protocol

### a) Validation of gentamicin susceptibility in *Staphylococcus aureus*

*Staphylococcus aureus* ATCC 27538 was cultured in the presence of increasing concentrations of gentamicin (0.5–8 µg/mL), and growth was monitored at 600 nm over 20 hours. Untreated controls exhibited typical exponential growth, reaching the stationary phase within 8–10 hours. Gentamicin exerted a dose-dependent inhibitory effect, partial inhibition with a prolonged lag phase and reduced optical density was observed at 0.5–2 µg/mL, whereas complete growth inhibition was achieved at concentrations ≥3 µg/mL. These results indicate that the minimum inhibitory concentration (MIC) lies between 2 and 3 µg/mL. The clear dose-response relationship, together with 12 biological replicates per condition, confirms the assay’s reliability and reproducibility (Fig. 11).

**Figure 11.**
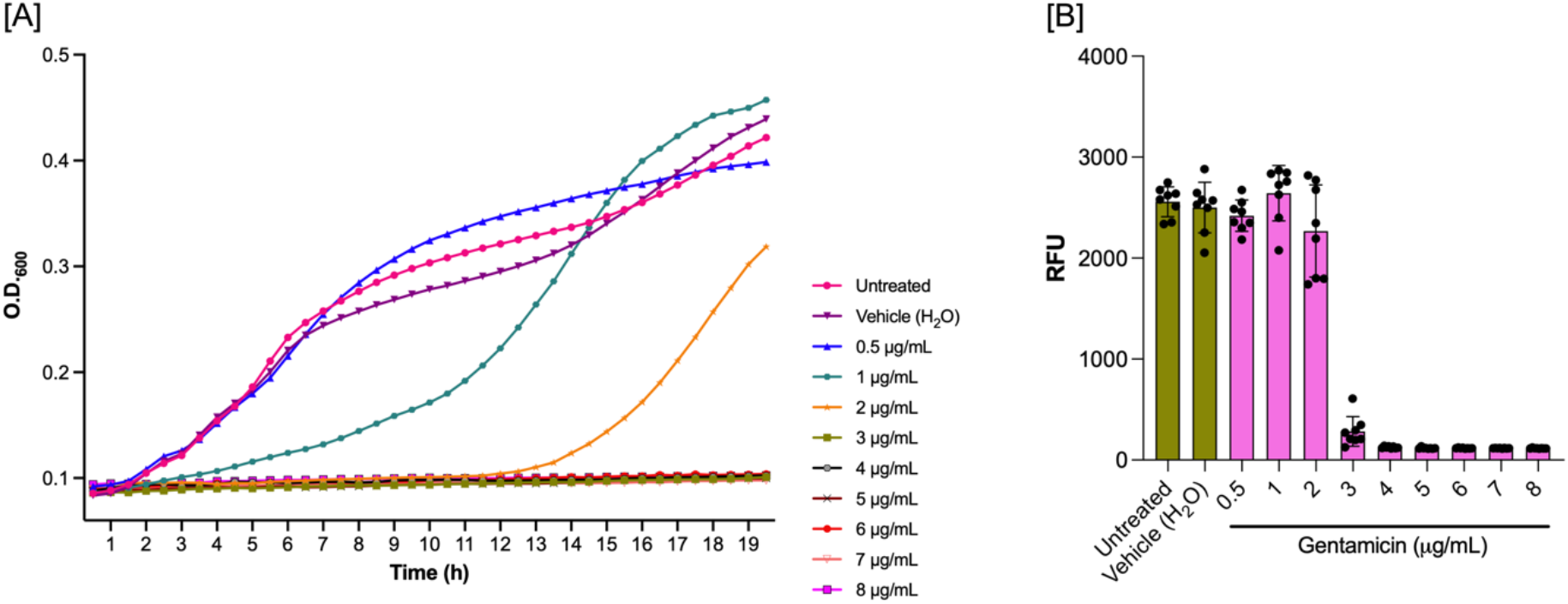
Validation of resazurin assay for assessing bacterial viability. (A) Growth kinetics of *Staphylococcus aureus* ATCC 27543 in culture medium over 20 hours. (B) Viability assessment using the resazurin reduction assay. Bacterial cultures were exposed to increasing concentrations of gentamicin (0.5–8 µg/mL) and monitored continuously for 20 hours. Resazurin reduction to resorufin (fluorescence) correlates with metabolically active cells. Data points represent mean values from nine biological replicates per condition.

### b) Assessment of actinomycin D cytotoxicity in 2D MDA-MB-231 cultures

MDA-MB-231 monolayers were treated with 10 µM actinomycin D for 24 hours, and morphological changes were assessed by bright-field microscopy (Fig. 12 A). Untreated and vehicle-treated cells (0.05% DMSO) displayed typical epithelial morphology with adherent, polygonal cells forming a confluent monolayer, confirming that DMSO is non-cytotoxic at this concentration. In contrast, cells exposed to 10 µM Actinomycin D exhibited pronounced cytotoxic effects, including cell rounding, detachment, reduced density, and floating debris. These findings, consistent across three independent experiments with triplicate samples, confirm the reproducibility of the 2D cytotoxicity assay (Fig. 12 B).

**Figure 12.**
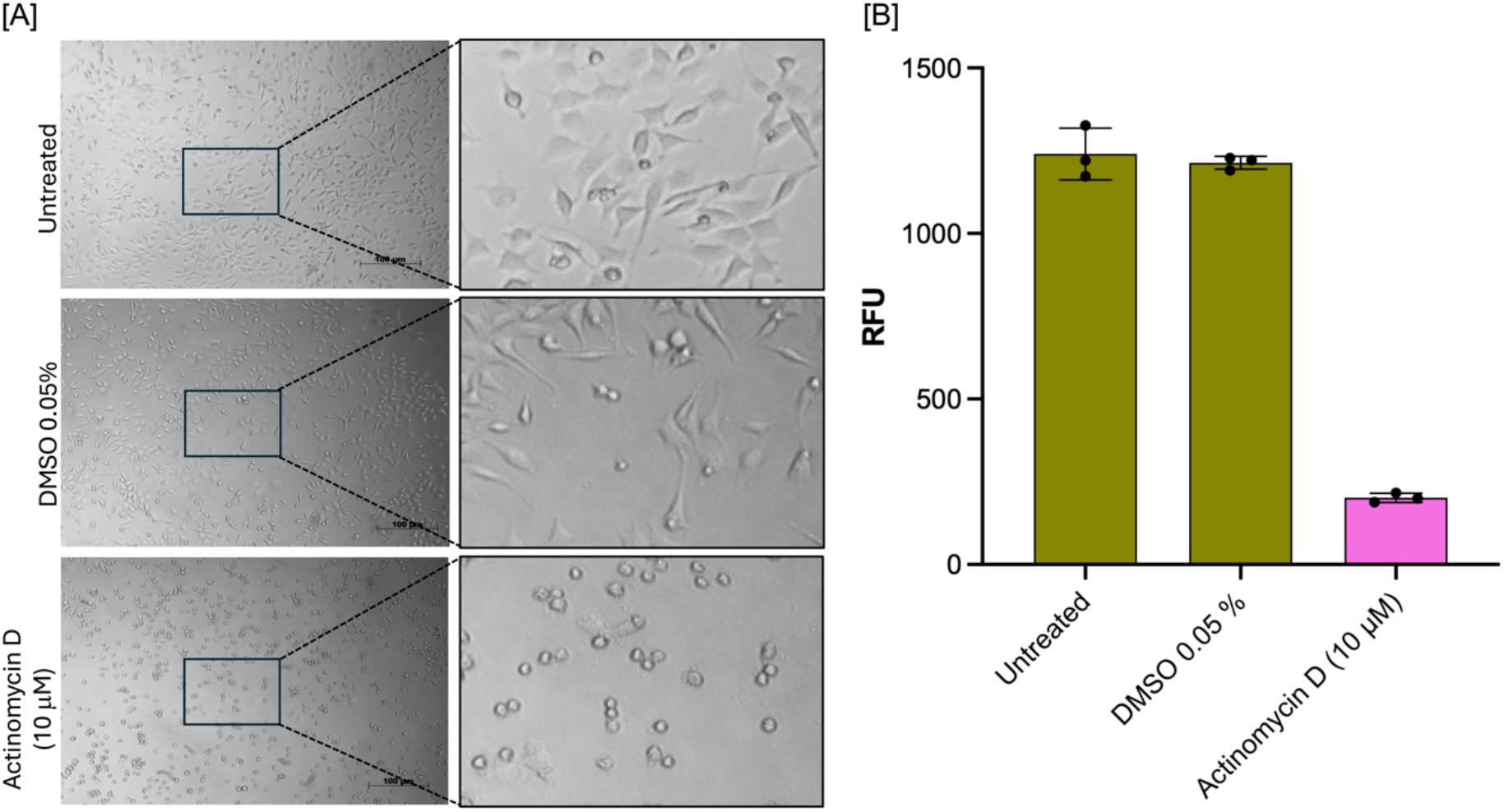
Validation of resazurin assay for assessing cytotoxicity in MDA-MB-231 breast cancer cells cultured in 2D monolayers. (A) Representative bright-field microscopy images (10× magnification) showing MDA-MB-231 cell morphology. (B) Quantitative viability assessment using the resazurin reduction assay. Cells were incubated for 24 hours under the following conditions: untreated control (incomplete medium); vehicle control (0.05% DMSO); and 10 µM Actinomycin D as a positive control for cytotoxicity. Resazurin reduction to fluorescent resorufin reflects metabolic activity of viable cells. Scale bar = 100 µm. Results are representative of three independent experiments, each performed in triplicate.

### c) Assessment of actinomycin D cytotoxicity assay in MDA-MB-231 spheroids

To assess the antiproliferative effect of actinomycin D in a three-dimensional context, MDA-MB-231 cells were first cultured under non-adherent conditions for 96 hours to allow formation of compact spheroids (Fig. 8). Following this establishment phase, spheroids were transferred to adherent plates and exposed to 10 µM actinomycin D for 24 hours. This experimental design enabled simultaneous observation of cell proliferation (via cells migrating from the spheroid and adhering to the plate) and cell death (via disruption of spheroid integrity).

Untreated control spheroids exhibited compact, three-dimensional architecture with smooth, well-defined borders. Upon transfer to adherent plates, these spheroids attached, and cells began to migrate outward, forming a proliferative halo around the core. Vehicle-treated spheroids (0.05% DMSO) showed behavior comparable to that of untreated spheroids, confirming that DMSO at this concentration does not affect spheroid integrity or proliferative capacity.

In contrast, spheroids treated with 10 µM Actinomycin D displayed marked disruption, including loss of compactness, irregular borders, loosening of cellular architecture, and the presence of dissociated cells surrounding the spheroid core. Notably, minimal cell migration or adherence was observed, indicating impaired proliferative capacity and significant drug-induced cytotoxicity (Fig. 13).

**Figure 13.**
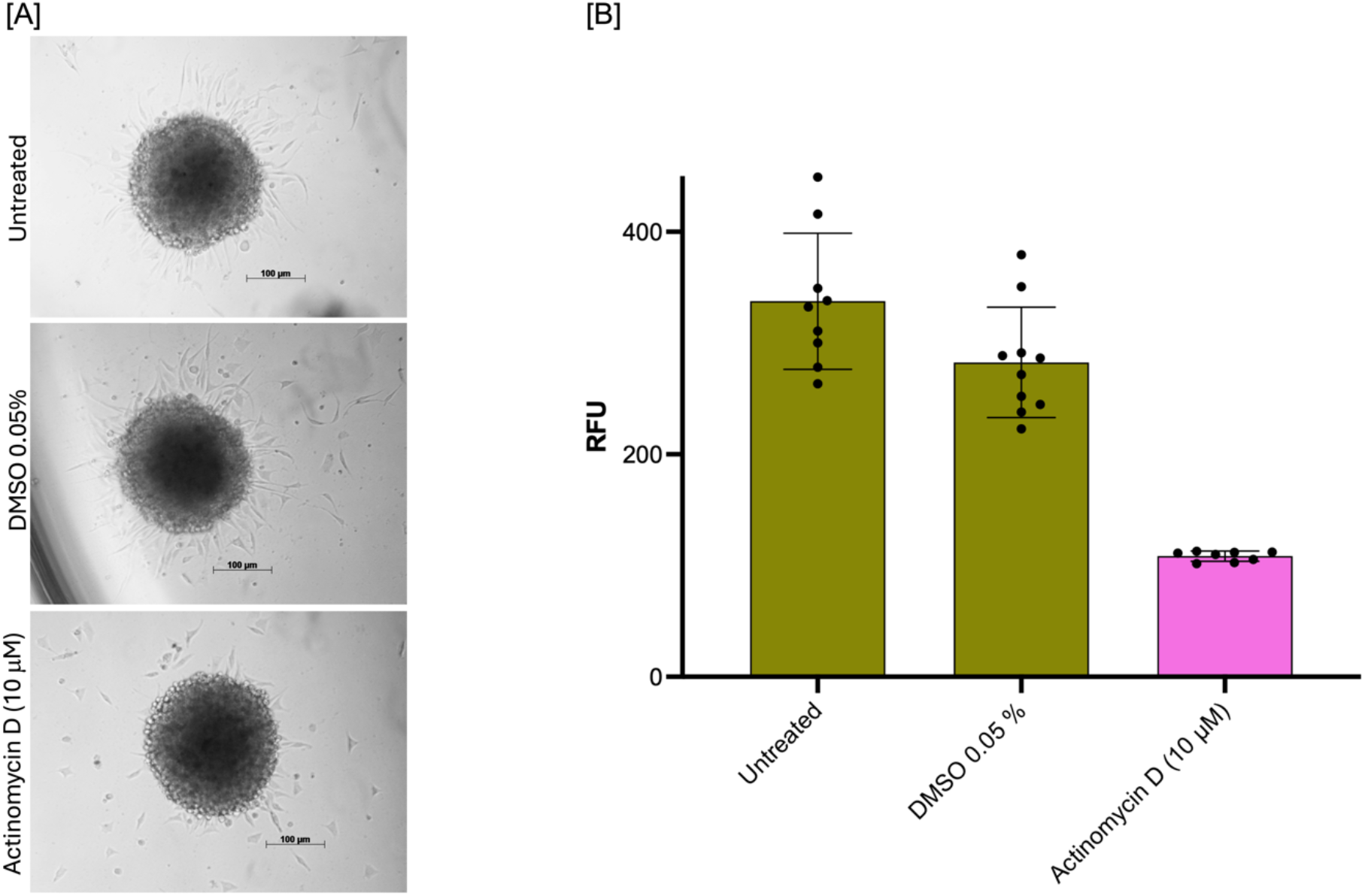
Validation of resazurin assay for assessing cytotoxicity in MDA-MB-231 breast cancer cells cultured as 3D spheroids. (A) Representative bright-field microscopy images (10× magnification) showing MDA-MB-231 spheroid morphology. (B) Quantitative viability assessment using the resazurin reduction assay. Cells were seeded into agarose-coated 96-well plates at a density of 20,000 cells per well and cultured under non-adherent conditions for 96 hours to allow formation of compact spheroids. Spheroids were then incubated for an additional 24 hours under the following conditions: untreated control (incomplete DMEM); vehicle control (0.05% DMSO); or 10 µM Actinomycin D (positive control for cytotoxicity). Resazurin reduction to fluorescent resorufin reflects the metabolic activity of viable cells. Scale bar = 100 µm. Images are representative of three independent experiments, each performed in triplicate.

In summary, the resazurin-based viability assay demonstrated remarkable versatility as a universal detection method, performing consistently across prokaryotic and eukaryotic systems and across different culture architectures. The protocols established herein are robust and reproducible for investigating antibacterial activity and cytotoxicity in both 2D cell culture monolayer and 3D spheroid cultures. This broad applicability positions resazurin as a valuable tool for diverse cell viability assessments in drug screening and mechanistic studies.

This protocol has been used and validated in the following research article(s):

**Cervantes-Rivera, R. *et al***. (Manuscript in preparation). Antitumor effects of avocado (*Persea americana* var. drymifolia) seed lipid extract on MDA-MB-231 cells in 2D and 3D models.

## General notes and troubleshooting

### A) General notes

| Aspect | Recommendation |
| --- | --- |
| **Sterility** | Maintain strict aseptic technique throughout all procedures. Perform all cell culture steps involving cell culture in a biosafety cabinet. |
| **Pre-warming** | Pre-warm all media, PBS, and trypsin solutions to 37 °C before use to avoid thermal shock to cells. |
| **Agarose coating** | Prepare agarose-coated plates fresh whenever possible. If storing, ensure they are well-sealed to prevent dehydration and contamination. |
| **Cell passage number** | Use cells between passages 3 after thawing for consistent spheroid formation and experimental reproducibility. |
| **Seeding density** | Optimize seeding density for each cell line. MDA-MB-231 cells typically form compact spheroids at 20,000 cells/well, but densities from 5,000 to 50,000 cells/well should be tested. |
| **Medium displacement** | When changing medium, always aspirate from the side of the well and add fresh medium slowly along the wall to avoid disturbing spheroids. |
| **Resazurin handling** | Protect the resazurin solution from light at all times. Prepare aliquots and store at –20 °C to avoid repeated freeze-thaw cycles. |
| **Controls** | Always include untreated controls, vehicle controls (e.g., DMSO at matching concentrations), and medium-only blanks on each plate. |

### B) Troubleshooting guide

| Problem | Possible cause | Solution |
| --- | --- | --- |
| **Low fluorescence signal** | Insufficient viable cells; resazurin incubation too short; spheroids too large for dye penetration. | Increase seeding density. Extend resazurin incubation time (try 6–8 hours). For large spheroids, consider lysing spheroids and measuring protein content or using a more penetrating viability dye. |
| **High background fluorescence** | Contamination; autofluorescence of medium or drug. | Check sterility. Include medium-only and drug-only controls. Use phenol red-free medium during the assay if necessary. |
| **Fluorescence varies widely between replicate wells** | Uneven spheroid size; pipetting error. | Ensure consistent spheroid formation. Use a multichannel pipette to add resazurin. Verify that spheroids are not lost during medium changes. |
| **Spheroids dislodge during resazurin addition** | Pipetting is too forceful. | Add resazurin solution very slowly along the well wall. Consider using a repeat pipettor for consistent, gentle dispensing. |
| **No color change observed (wells remain blue)** | All cells are dead; resazurin degraded. | Check negative control (untreated spheroids) to verify assay functionality. Ensure resazurin stock is stored in a protected, light-free location and is not expired. |
| **The microplate reader detects bubbles** | Bubbles formed during pipetting. | Tap the plate gently to dislodge bubbles before reading. Use a needle to pop any persistent bubbles. Centrifuge plate briefly if necessary (but avoid disturbing spheroids). |
| **High well-to-well variability in fluorescence** | Spheroids not centered; meniscus effect. | Ensure spheroids are at the bottom center. Some readers have reduced sensitivity at edge wells; consider using only inner wells for experiments. |
| **Fluorescence saturation** | Signal too high; gain setting incorrect. | Reduce resazurin incubation time. Dilute samples before reading. Adjust reader gain settings. |

### C) Quick reference checklist for troubleshooting

| Observation | First actions to try |
| --- | --- |
| **Poor spheroid formation** | Increase seeding density, optimize shaking, and check agarose coating. |
| **Spheroids lost during medium change** | Aspirate/add along wall, use gentle pipetting, leave residual volume. |
| **No drug effect** | Increase concentration, extend treatment time, and verify drug stability. |
| **Low-resazurin signal** | Extend incubation time, increase seeding density, check dye integrity. |
| **High variability** | Standardize technique, use a multichannel pipette, and verify cell counting. |
| **Contamination** | Review aseptic technique, discard contaminated plates, prepare fresh media. |

## Acknowledgment

We gratefully acknowledge funding from Coordinación de la Investigación Científica de la UMSNH (Proyecto 17644). We are grateful for postdoctoral stipend support to Ramón Cervantes-Rivera from Secretaría de Ciencia, Humanidades, Tecnología e Innovación (SECIHTI).

## Author contribution

**Conceptualization:** Ramón Cervantes-Rivera

**Methodology:** Ramón Cervantes-Rivera

**Investigation:** Ramón Cervantes-Rivera, Atalia Ziret Romero Rosas, Sandra Jetsamari Figueroa Ortíz, Luisa Nirvana González-Fernández

**Formal analysis:** Ramón Cervantes-Rivera, Atalia Ziret Romero Rosas, Sandra Jetsamari Figueroa Ortíz

**Data curation:** Ramón Cervantes-Rivera

**Writing – Original draft:** Ramón Cervantes-Rivera, Atalia Ziret Romero Rosas

**Writing – Review & Editing:** Ramón Cervantes-Rivera, Atalia Ziret Romero Rosas, Luisa Nirvana González-Fernández, Sandra Jetsamari Figueroa Ortíz, Alejandra Ochoa Zarzosa and Joel E. López-Meza

**Visualization:** Ramón Cervantes-Rivera, Luisa Nirvana González-Fernández, Sandra Jetsamari Figueroa Ortíz

**Funding acquisition:** Joel E. López-Meza and Alejandra Ochoa Zarzosa

**Supervision:** Ramón Cervantes-Rivera, Alejandra Ochoa Zarzosa and Joel E. López-Meza

**Project administration:** Ramón Cervantes-Rivera and Joel E. López-Meza

